# Pollen Profiles and Floral Sources of Tennessee Honeys, an Influence of Appalachian Ecoregions

**DOI:** 10.1101/2025.06.27.660778

**Authors:** Conner R. Reed, Larry J. Millet

**Affiliations:** School of Forest, Fisheries, & Geomatics Sciences, University of Florida. Gainesville, FL 32611-0410; Department of Civil and Environmental Engineering, University of Tennessee, Knoxville. Knoxville, TN 37996

## Abstract

Tennessee and the surrounding Appalachian region offer a uniquely rich floral landscape that supports the production of many flavorful, high-quality types of honey. Plant species such as tulip poplar, black locust, basswood, sumac, and sourwood provide abundant nectar that supports strong honey yields. Pollen from plants that produce little or no nectar also contribute to a diversity of honey flavors. This study uses DNA-based pollen profiling to identify plant sources contributing to honey varieties in single frames of honey. By analyzing individual honey frames through molecular and bioinformatic techniques, the project supports how regional and local diverse plant communities contribute to honey composition.

This work demonstrates how genomic tools can be applied to classify honey by floral source, supporting future research in molecular genomics, apiculture, and biodiversity mapping. The findings also inform efforts to promote small batch honey extraction to enhance consumer awareness and valuation of locally sourced honey products.

## Introduction

Tennessee and neighboring Appalachian beekeepers are uniquely positioned to produce and market some of the most flavorful and distinctive varieties of honey in the Continental United States. Identifying the pollen sources and floral profiles in Tennessee and Appalachian honey has the potential to play a crucial role in supporting quality honey producers who operate in a challenging honey market that contains imported honey. With better tools and data to understand and define their honey, beekeepers can educate consumers and showcase the exceptional variety of flavors that come from Tennessee’s diverse landscapes. This pollen profiling project is intended to aid informing regional beekeepers and promote broader public understanding of the ecological and botanical factors that influence honey composition, highlighting the role of southern Appalachian^1,2^ plant biodiversity and the meaningful contributions of small-scale apicultural practices.

Despite the widespread presence of low-cost, mass-produced honey blends in grocery stores, consumers consistently recognize the superior quality of authentic, locally-produced honey often provided by hobbyists who have 1-10 hives and sideline beekeepers who manage 11-50 hives. These small-batch specialty crop honey products can preserve the floral character of the blooms that feed the bees, which offers a unique tasting experience appreciated by customers who are interested in pollen’s nutritional benefits, possible allergenic relief, pollinator health, honey production, and regional agriculture.

The Appalachian region hosts an exceptionally rich diversity of flowering plants due to its complex topography, varied microclimates, and ancient geological history. This biodiversity includes a wide array of native trees, shrubs, and ground flowers that bloom from early spring through late summer and provide consolidated foraging periods for pollinators.^3^ In the East Tennessee region, most of the important nectar-producing plants for honey production are trees. Tree blossoms play a central role in sustaining both wild and managed bee populations across the region.

In the Tennessee portion of Appalachia, flowering plants are especially pronounced in the Central Ridge and Valley, and Blue Ridge Mountain ecoregions, where both upland hardwoods and mixed mesophytic forests thrive. Figure 1 summarizes the regional descriptions for the Interior Low Plateau-Highland Rim, Northern Cumberland Plateau, Northern Cumberland Mountains, and the Central Ridge and Valley in Tennessee from Nashville, to the Blue Ridge Mountains in Asheville, North Carolina. In this region, many of the dominant deciduous trees are oak and hickory, which are wind-pollinated and not significant nectar sources for honey bees. Other deciduous species are also present, particularly in the Central Ridge and Valley; these deciduous species do produce nectar and are of interest to pollinators, such as honey bees.

**Figure 1.**
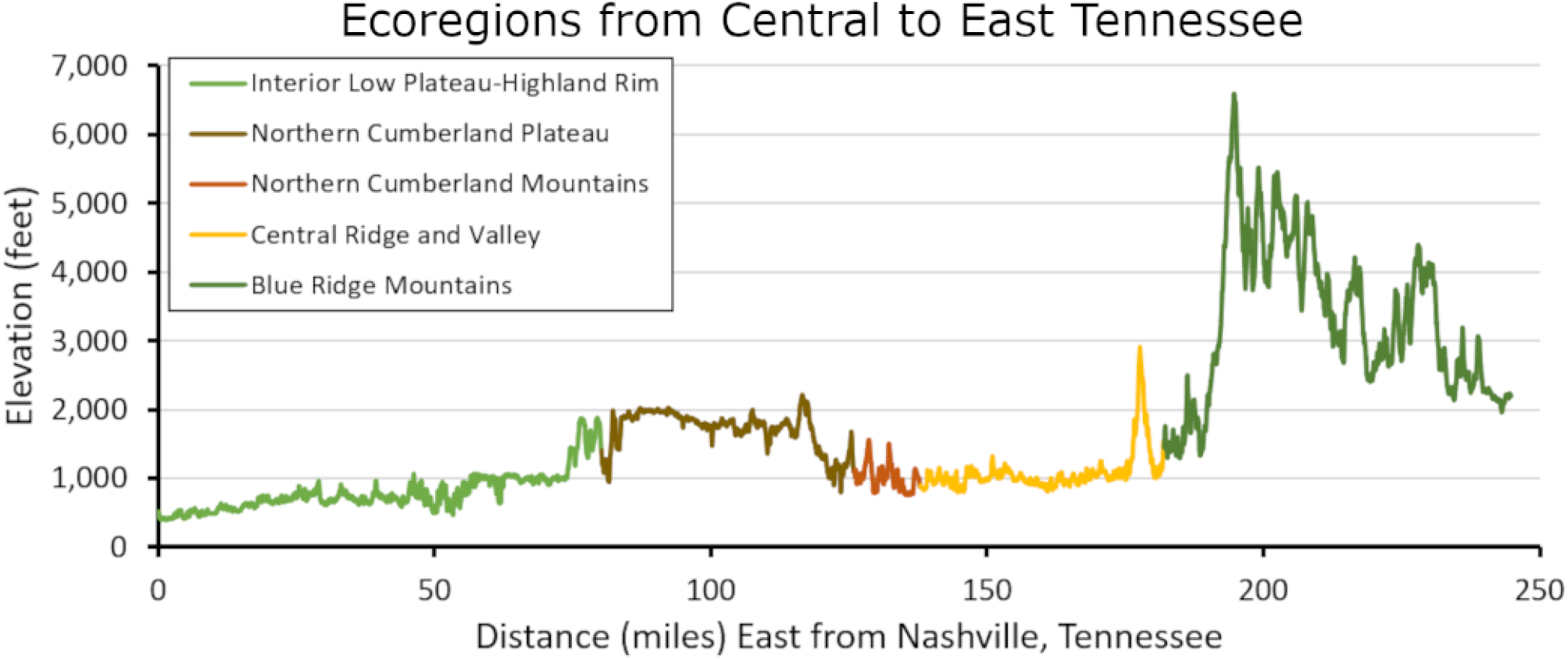
A topographic profile of the East Tennessee region. The graph highlights the elevation and ecoregions in an east-to-west elevation profile^5^ connecting points from Nashville to Knoxville, Tennessee to the highest point in the Great Smokey Mountains National Park, and ending in Asheville, North Carolina. Ecoregions are classified according to the National Hierarchical Framework of Ecological Units. ^6-7^

The maple (genus, *Acer*), pear (*Pyrus*) and black cherry (*Prunus serotina*) trees provide an essential early spring supply of nectar and pollen, supporting the vital growth of bee colonies during post-winter buildup. Specifically, pear tree pollen has been identified as one of the highest nutritional value pollen types, containing significantly greater levels of five essential amino acids— isoleucine, phenylalanine, methionine, histidine, and lysine—compared to other pollens. Consequently, pear trees may better support honey bee health and physiological function.^4^

The tulip poplar (*Liriodendron tulipifera*) and black locust (*Robinia pseudoacacia*, in the pea family) stand out as top nectar sources for honey bees throughout the Eastern U.S., notably contributing to the popularity of tulip poplar honey found in Tennessee. The American basswood (*Tilia americana*) native to Tennessee, blooms in early summer and offers abundant nectar that produces a light, fragrant honey highly prized by beekeepers. Sourwood or sorrel tree (*Oxydendrum arboreum*), a standout native species, blooms in midsummer and is renowned for producing a prized honey highly sought after by both beekeepers and consumers. Together, these trees play a crucial role in sustaining honey bee populations in forested landscapes.

### Interior Low Plateau and Highland Rim (400 to 1,900 feet)

This region features oak-hickory forests, cedar glades, and scattered prairies. Dry sites support post, southern red, scarlet, chestnut, and blackjack oaks; moist areas favor white and black oaks. Hickories like pignut and shagbark are common. Black locust and black cherry—both important nectar sources—occur along edges and disturbed areas. Much of the natural vegetation of this area has been cleared for agriculture.

### Northern Cumberland Plateau (850 to 2,300 feet)

This part of Tennessee hosts a mix of Appalachian oak and mesophytic forests. In Kentucky, shortleaf pine and oak dominate; in Tennessee, oaks are more common while pines are limited. Hickory remains a minor portion of the tree population. Tulip poplar and black locust appear in disturbed or early successional areas, providing strong spring nectar flows. Roughly 20% of this ecoregion has been cleared for agriculture.

### Northern Cumberland Mountains (760 to 1,530 feet)

These forests include Appalachian oak, mesophytic hardwoods (tulip poplar, red maple, basswood), and northern hardwoods at higher elevations. Common oaks include white, black, scarlet, and blackjack; hickories such as mockernut and pignut are present. Tulip poplar, basswood, and black locust offer substantial nectar during spring and early summer. Most of this area has been cleared of natural vegetation for agricultural use. Strip mining for coal has disturbed about 5 percent of the area.

### Central Ridge and Valley (800 to 2,900 feet)

These locations support oak-pine forests, with widespread composition of shortleaf pine trees. Oak species shift with soil moisture: drier slopes favor post, scarlet, chestnut, and blackjack oaks; wetter areas support white and southern red oaks. Deciduous nectar sources include sweetgum, blackgum, tulip poplar, and black locust. In the south, loblolly pine becomes more common. Over 60% of the landscape is agricultural or pasture.

### Blue Ridge Mountains (1,293 to 6,590 feet)

The Great Smoky Mountains are a subrange of the Appalachian Mountains, which form part of the Blue Ridge Physiographic Province, these mountains host diverse plant communities: dry slopes support oak species; valleys favor mesophytic trees like tulip poplar, basswood, and red maple; higher elevations transition to red spruce and Fraser fir, Black locust, eastern white pine, and Table Mountain Pine (*Pinus pungens* is a small pine native to the Appalachian Mountains) appear in fire-adapted zones and disturbed areas. Roughly 35% has been cleared, primarily in broad valleys.

Within each ecoregion, numerous subregional variations arise from a range of factors, including: underlying geological differences that influence the location of cities, towns, farms, and forested areas; the composition of tree species that are supported by specific geological formations; patterns of tree removal and replacement associated with forestry practices; and prevailing tree and landscaping preferences that occur both within and between urban and rural communities.

## Single-frame honey profiles

Variation in plant species across ecoregions and subregions contributes to the diversity of honey varietals available for production. To better understand and characterize the range of honey products generated in central and eastern Tennessee, we conducted single-frame honey extractions using the crushed comb technique. Honey frames were obtained from a hive located at the University of Tennessee Forest Resources AgResearch and Education Center (FRAREC), Oak Ridge, Tennessee. This method was selected to circumvent methodological challenges associated with using centrifugal extractors for single-frame honey collection. Figure 2 shows an image of a portion of a single frame of honey that displays an example of the range of honey colors that can be produced in East Tennessee; in particular, it demonstrates how honey bees can store different honey types within a single frame. This is a familiar sight to beekeepers, but not an experience appreciated by the public. Figure 3 shows representative graphs of different single-frame honey produced in our hives kept at the University of Tennessee FRAREC, Oak Ridge, Tennessee, which does not represent all aspects of East Tennessee and Appalachian subregions and microclimates; however, it is a suitable location to test frame-by-frame pollen mapping through pollen DNA sequencing of honey varieties.

**Figure 2.**
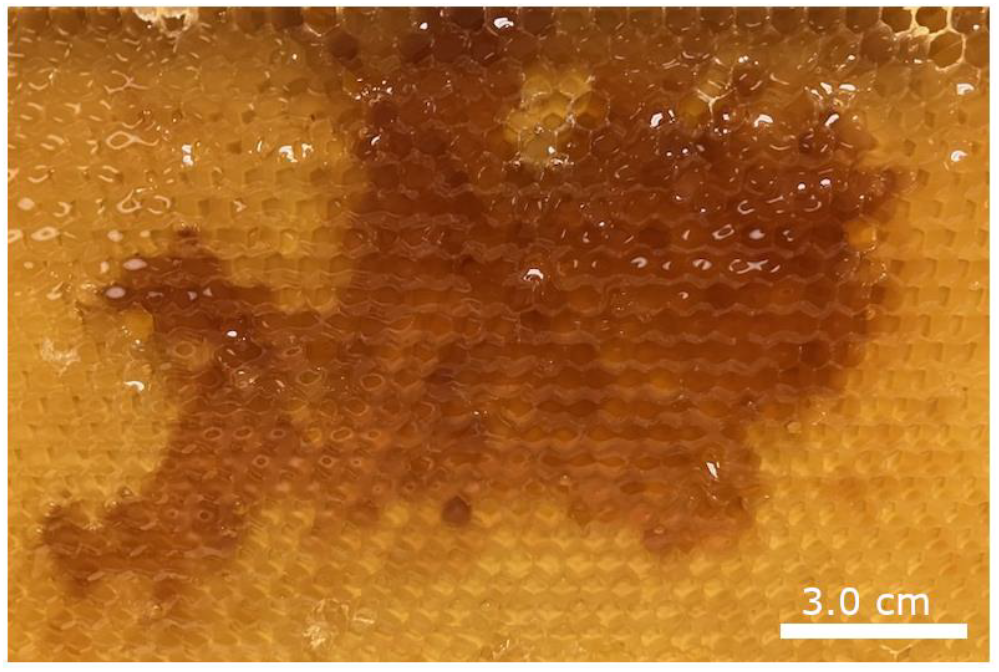
Throughout the three-month spring nectar flow in East Tennessee, many varieties of light and dark honey are produced. This subsection of a single honey frame shows the variations that can be observed and collected.

**Figure 3.**
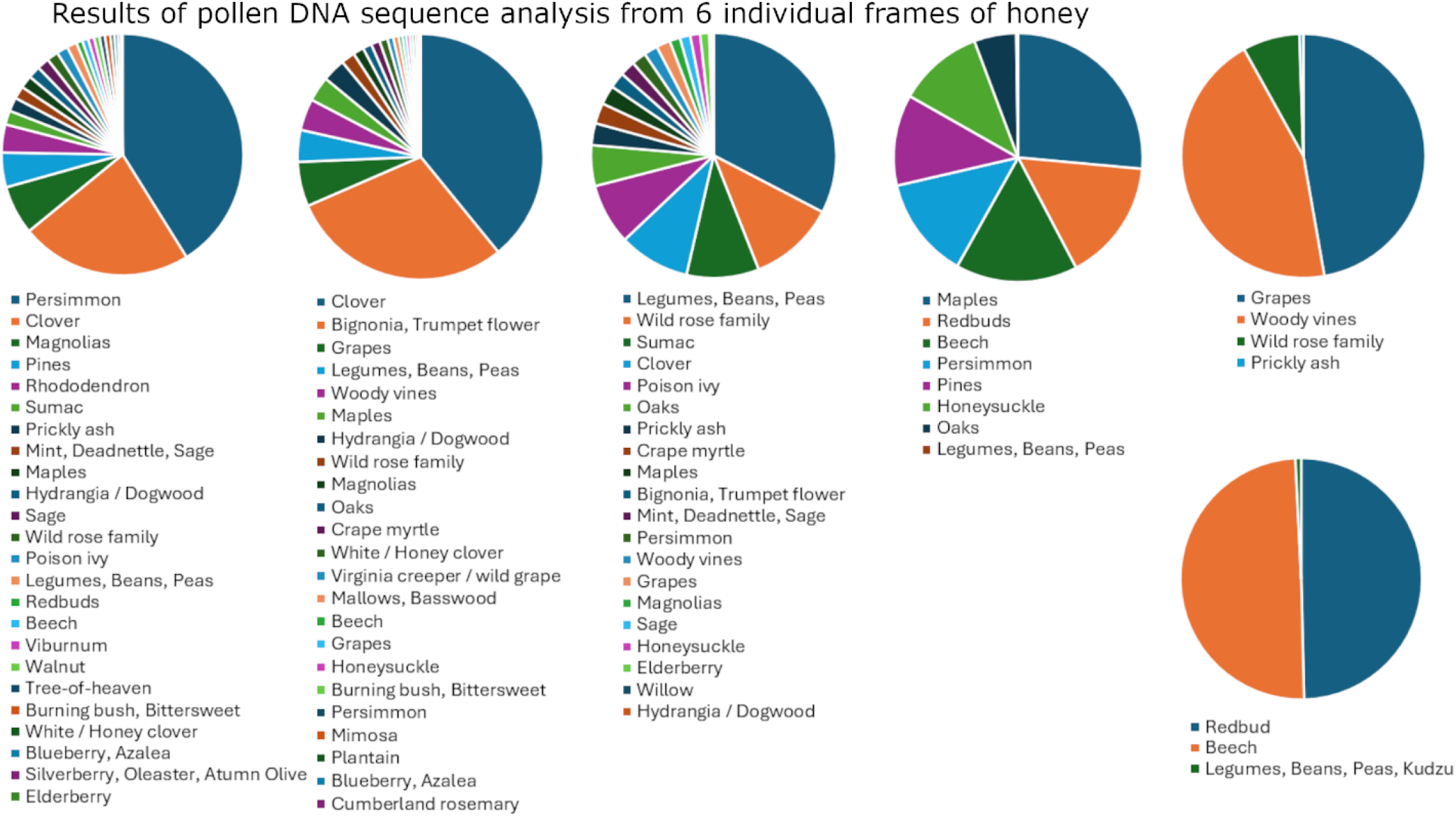
Proportional representation of pollen categories from DNA sequence reads obtained from six frames of honey (crushed comb sampling). Each pie chart contains common names listed from most abundant (top) to least abundant (bottom). Honey frames were harvested at the research apiary within the University of Tennessee Forest Resources AgResearch and Education Center, Oak Ridge, Tennessee; Ecoregion – Central Ridge and Valley.

## Pollen DNA sequence methods

Samples were analyzed through pollen DNA analysis (Jonah Ventures, LLC). Pollen was extracted from raw honey by diluting 10 mL of honey in 40 mL of sterile water, followed by heating at 65°C for 20 minutes with periodic mixing. After centrifugation at 4,000 RPM for 30 minutes, the resulting pollen pellet was collected, swabbed, and transferred into extraction vessels. Genomic DNA was extracted using the DNeasy 96 PowerSoil Pro Kit (Qiagen, Cat# 47017) and eluted in 100 µL. The chloroplast trnL intron was PCR-amplified from each sample using c and h trnL primers with Illumina adapter overhangs,^8^ followed by gel verification and enzymatic cleanup. A second, indexing PCR appended unique 12-nucleotide barcodes to each sample. Final libraries were normalized using SequalPrep plates (Life Technologies, Cat# A10510-01), pooled, and sequenced on an Illumina MiSeq (v2 500-cycle kit). Bioinformatic processing included demultiplexing with Pheniqs, primer trimming with Cutadapt, read merging and quality filtering with VSEARCH, and denoising using UNOISE3. Exact sequence variants (ESVs) were assigned taxonomies via a best-hit algorithm against a custom database, incorporating GenBank and Jonah Ventures plant barcode records. This entire DNA sequencing process is commercially available and accessible to the range of small-to large-scale beekeepers.

Pollen source identities are presented using common names to enhance accessibility and relevance for the beekeeping community and the general public. The samples are arranged in order of complexity where a high number of pollen varieties detected in the honey are evident, juxtaposed to other single-frame honey varieties of relative simplicity. We look at the few of the frames to gain a greater understanding of pollen content in honey production and see complex representation of pollens even among samples collected at a single location.

In East Tennessee, maple trees generally bloom earlier than, but often overlapping with, the eastern redbud (*Cercis canadensis*) in late winter and early spring. Maple and redbud typically bloom from late February and mid-March, with the latter extending into early April, depending on the ecoregion. They thrive along forest edges and roadsides and are highly attractive to bees, offering an important early-season source of both nectar and pollen. In contrast, American beech (*Fagus grandifolia*) blooms about 2–4 weeks later, from mid-to late April, and is wind-pollinated with low nectar value to pollinators, but it is an available source of pollen. Beech trees are more common in moist, shaded upland forests.

Grapevines (*Vitis spp*.) bloom from late May to early June, often climbing along edges and open woodlands. Their small flowers offer a moderate nectar and pollen source for insects. Grapes are part of the woody climbing vines (Order: *Vitales*, Family: *Vitaceae*) that can bloom slightly later; from late May through July depending on species and conditions; they occur in disturbed or shaded habitats and may offer low to moderate floral value to pollinators during early summer.

These species represent a subset of the common trees and plants identified through DNA-based pollen analysis of honey samples. The bloom succession of key nectar-producing tree species utilized by honey bees in the East and Central Tennessee ecoregions follows a general seasonal pattern each year:

Maple → Cherry → Black locust → Tulip popular → Privet → Sumac → Sourwood.

Annual variation in weather conditions can significantly influence the timing and duration of floral blooms, while periods of rainfall may substantially reduce or entirely suppress the availability of nectar and pollen from flowering plants during a given period of days or weeks.

## Conclusion

East Tennessee’s rich biodiversity and native flowering trees—like tulip poplar, basswood, black locust, and sourwood—support the production of exceptional, locally distinctive honeys. Through pollen DNA analysis of honey, we gain a deep appreciation of the rich variety of pollen types contained within a single frame of honey. These examples demonstrate, in a preliminary and simplified form, the variety and complexity of pollen content that can occur in honey from East Tennessee—and, by extension, from the broader Appalachian region. For the public, it underscores the importance of supporting local beekeepers, protecting native plant communities that sustain pollinators, and contributing to the region’s agricultural identity.

## Acknowledgements

We are grateful to the many beekeepers of Tennessee for their comments, efforts, and interest in this work. The staff of the University of Tennessee Forest Resources Research and Education Center (the UT Arboretum) for making this work possible. C. Foster, C. Lowden, S. Greenwood for manuscript review and edits. Jonah Ventures performed pollen DNA processing and sequencing. Funding for *Improving Quality Production and Marketing of Authentic Tennessee Artisanal Honey in a Deteriorating Honey Market* was made possible by the U.S. Department of Agriculture’s (USDA) Agriculture through grant AM22SCBPTN1113. Its contents are solely the responsibility of the authors and do not necessarily represent the official views of the USDA.

